# A brain-wide analysis maps structural evolution to distinct anatomical modules

**DOI:** 10.1101/2022.03.17.484801

**Authors:** Robert A. Kozol, Andrew J. Conith, Anders Yuiska, Alexia Cree-Newman, Bernadeth Tolentino, Kasey Banesh, Alexandra Paz, Evan Lloyd, Johanna E. Kowalko, Alex C. Keene, R. Craig Albertson, Erik R. Duboue

## Abstract

Brain anatomy is highly variable and it is widely accepted that anatomical variation impacts brain function and ultimately behavior. The structural complexity of the brain, including differences in volume and shape, presents an enormous barrier to define how variability underlies differences in function. In this study, we sought to investigate the evolution of brain anatomy in relation to brain region volume and shape across the brain of a single species with variable genetic and anatomical morphs. We generated a high-resolution brain atlas for the blind Mexican cavefish and coupled the atlas with automated computational tools to directly assess brain region shape and volume variability across all populations. We measured the volume and shape of every neuroanatomical region of the brain and assess correlations between anatomical regions in surface, cavefish and surface to cave F_2_ hybrids, whose phenotypes span the range of surface to cave. We find that dorsal regions of the brain are contracted in cavefish, while ventral regions have expanded. Interestingly, in hybrid fish the volume and shape of dorsal regions are inversely proportional to ventral regions. This trend is true for both volume and shape, suggesting that these two parameters share developmental mechanisms necessary for remodeling the entire brain. Given the high conservation of brain anatomy and function among vertebrate species, we expect these data to studies reveal generalized principles of brain evolution and show that *Astyanax* provides a system for functionally determining basic principles of brain evolution by utilizing the independent genetic diversity of different morphs, to test how genes influence early patterning events to drive brain-wide anatomical evolution.

## Introduction

Brain anatomy is highly variable across the animal kingdom, yet little is known about the general evolutionary principles driving anatomical evolution of the brain (1-5). Comparative studies have provided insight into evolutionary and developmental mechanisms influencing anatomical change, but a lack of direct genetic and functional experiments across species have remained a major impediment. Two central hypothesis are thought to drive anatomical brain evolution; the developmental constraint hypothesis, that brain regions change together in a concerted matter, with selection operating on developmental mechanisms that govern the growth of all regions, and the functional constraint hypothesis, positing that selection can act on individual brain regions, and that regions which are functionally related will anatomically evolve together independent of other brain regions (6-8). While data supporting each hypothesis exist, the large divergence times and poor understanding of evolutionary history in most comparative models makes generalizing these theories difficult. Moreover, the relationship between the evolution of the size and shape of distinct anatomical regions is poorly understood, and it is unclear how these two important aspects of neuroanatomy explain the evolution of the brain.

Volume and shape govern anatomical variation across the brain and are thought to involve both overlapping and distinct mechanisms (9). However, most comparative studies tend to focus on either volume or shape, with some volume to shape analyses comparing trends across independent studies (10-12). Current models to explore mechanisms driving volume and shape, rely on non-model systems that lack experimental approaches, or model organisms that lack genetic diversity, creating an impediment for investigating basic principles of brain evolution. (13-18). A major improvement in our understanding of the anatomical evolution of brain shape and volume involves utilizing an evolutionary model that provides laboratory interrogation with high genetic variability.

The blind Mexican cavefish *Astyanax mexicanus* provides a powerful model for directly testing how genetic variation underlies brain-wide anatomical evolution (19, 20). *A. mexicanus* exists as a species with two robustly distinct forms: river dwelling surface fish and cave dwelling populations that have independently evolved troglobitic phenotypes (21, 22). This separation has led to high genetic diversity, that is unique to each population and underlies the stark differences in phenotypes between surface and cave populations (23, 24). Importantly, surface to cave hybrid offspring are biologically viable, allowing us to exploit the unique genetic diversity of each population, by relating differences in neurological features among hybrid individuals to genetic variants found in wildtype parents (25, 26). Therefore, these tools can be used to study covariation of neuroanatomy across a well-annotated atlas, determining the relationship between brain region shape and volume, which will be critical in understanding how brain regions evolve in relationship to one another.

In the current study, we generated a brain-wide neuroanatomical atlas for all *Astyanax* morphs and applied new computational tools for assessing brain-wide changes in both brain region volume (27) and shape (28). We then used this atlas and applied it to hybrid brains to make functional associations between naturally occurring genetic variation of wildtype populations and neuroanatomical phenotypes. Our data reveal that both brain-region volume and shape are impacted genetically brain-wide to influence the anatomical evolution of cavefish brains. Volume and shape variation brain-wide share developmental mechanisms that are causing cavefish brains to contract dorsally and expand ventrally. These results suggest that selection may be operating on simple developmental mechanisms, that likely impact early patterning events to modulate the volume and shape of brain regions.

## Results

### Generation of a single brain-wide atlas for all Astyanax morphs

To analyze regional variation in brain anatomy, we created a single atlas for all *Astyanax* morphs to provide neuroanatomical comparisons across surface, cave, and surface to cave hybrid populations. A neuroanatomical analysis pipeline from zebrafish that performs automated segmentation of brains was adapted and tested on *Astyanax* brains (27) (Figure1a&b and Supplementary Data Figure 1a-c). This tool provides a single atlas that can be continually segmented through brain regions to identify neuroanatomical differences across various molecularly and functionally defined sub-nuclei. (Figure 1c, Supplementary Data, Supplementary Figure 1d, Supplementary Tables 1-6). The segmentation accuracy was confirmed by comparing manual and automated segmentation (Supplementary Data, Supplementary Figure 2a&b, Supplementary Tables 7-10). We further confirmed these findings using molecular markers, such as the *insulin gene enhancer protein ISL-1 and Insulin gene enhancer protein ISL-2* (*Islet1/2*) and pairwise comparisons (Supplementary Data, Supplementary Tables 11-16). Volumetric and shape data were then measured and analyzed for variation across surface and cave brains (Supplementary Data, Supplementary Figure 1e and Figure 3a&b, Supplementary Tables 17-32). The atlas was then applied to the Pachón cavefish, Rio Choy surface fish and Pachòn to Rio Choy hybrid brains immunostained for anatomical structure, allowing us to address the evolutionary mechanisms underlying adaptations in brain anatomy.

**Figure 1.**
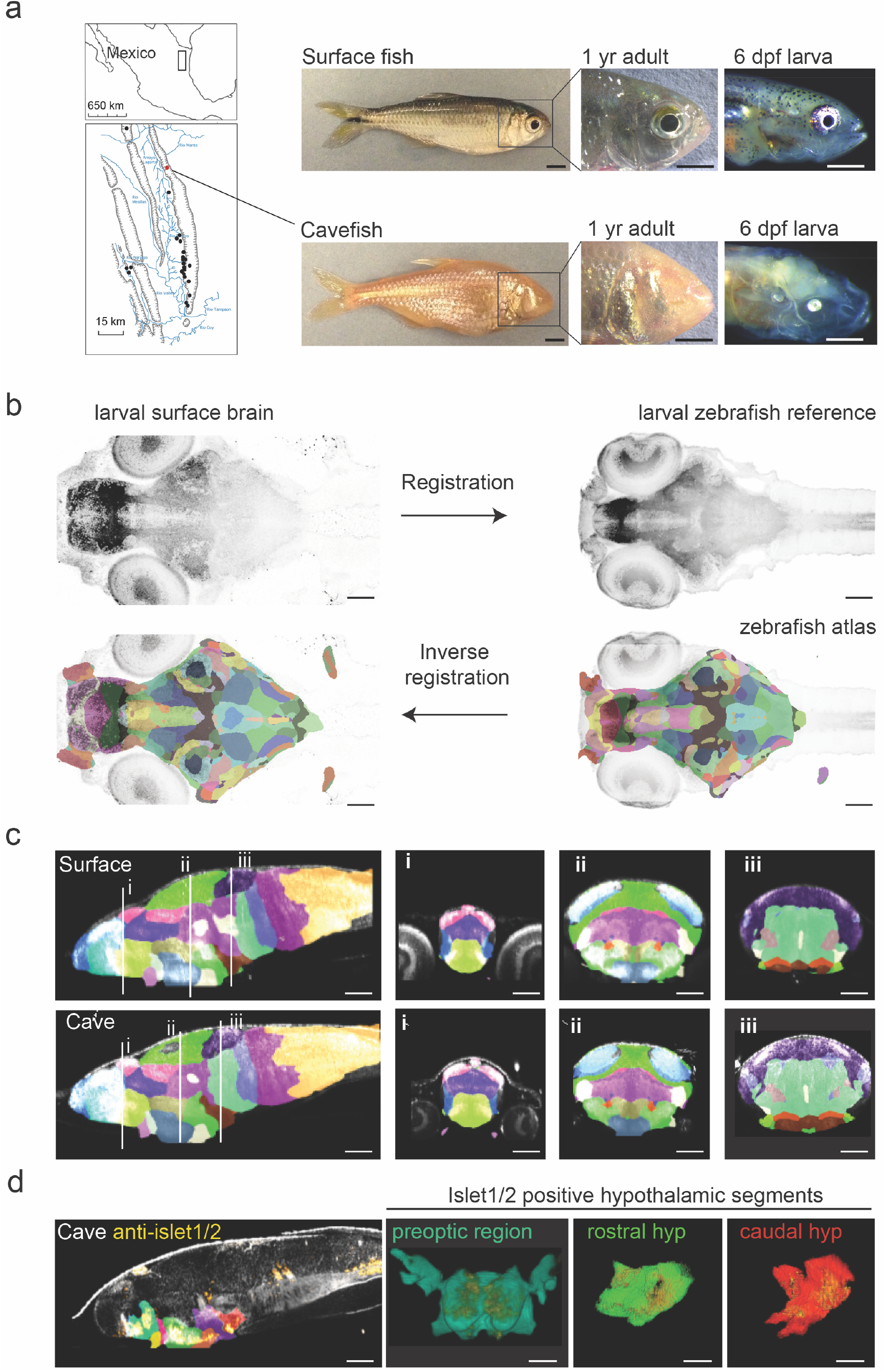
Developing a single *A. mexicanus* atlas to perform direct brain-wide morphometric analyses across all populations. (**a**) Map showing the 29 independently evolved cave populations (black dots) of the el Abra region in Mexico. The Pachón cavefish population used for this project is marked as a red dot. Scale bars = 0.5 cm (full fish, 1 yr adult) and 0.5 mm (larvae) (**b**) Schematic showing registration and atlas inverse registration method used to create an *A. mexicanus* atlas for cross population segmentation and analysis. (**c**) Sagittal and transverse (**i-iii**) sections of the 26 region surface fish and cavefish atlas. (**i**.) Habenula (pink), pallium (blue), ventral thalamus (purple) and preoptic (light green). (**ii**.) optic tectum neuropil (sky blue), optic tectum cell bodies (green), tegmentum (light purple), rostral hypothalamus (dark blue), posterior tuberculum (gold), statoacoustic ganglion (beige). (**iii**.) Cerebellum (dark purple), prepontine (light green), locus coeruleus (brown), raphe (beige), intermediate hypothalamus (dark brown) and caudal hypothalamus (bright red) (**d)** Islet1/2 antibody segmentation following ANTs inverse registration of cavefish atlas. Islet positive neurons fall within regions that have been reported islet positive in zebrafish. Scale bars (b-d) = 80 μm.

### Determining neuroanatomical variation brain-wide for surface and cave populations

To determine volumetric variation in an unbiased way, we compared regional variation across different levels of segmentation within our *A mexicanus* brain atlas. Progressively segmenting brains through sub-nuclei provided an analytical tool for defining regional variability, with localization of variability increasing as we scaled through each atlas (Figure 2a-c, Supplementary Tables 33-40; Supplementary Data Figure 3a&b, Supplementary Tables 17-32). While this comparison found volumetric differences that were previously reported in broad developmental regions (29, 30), such as the hypothalamus (Figure 2b, Supplementary Table 34), our atlas was able to isolate these differences to the intermediate and caudal hypothalamus (Figure 2c, Supplementary Tables 39-40). In addition, we also discovered novel volumetric differences, including contraction of the dorsal diencephalon in cavefish (Figure 2b, Supplementary Table 33), that we localized to the dorsal thalamus (Figure 2c, Supplementary Table 37). Overall, we were able to use this single atlas to pinpoint discrete differences between brain regions of surface fish and cavefish, along with a brain-wide model for how anatomical structure has been remodeled in cavefish (Figure 2d).

**Figure 2.**
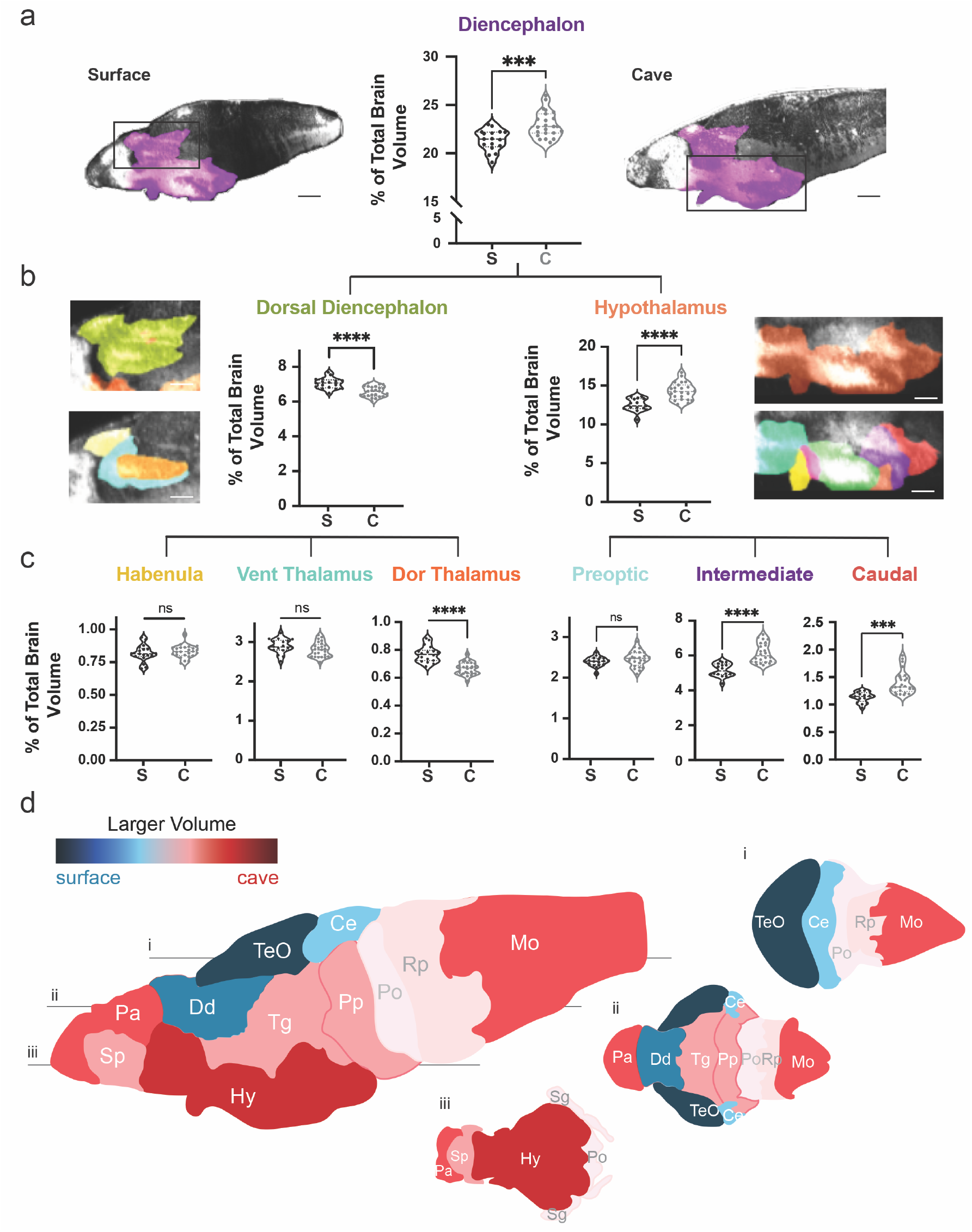
Volumetric variation in wildtype populations reveal regional contraction and expansion across cavefish brains. (**a)** Volumetric comparison of the diencephalon in surface and Pachòn cavefish. Percent total brain volume represents pixels of segment divided by total pixels in the brain. Sagittal sections show major brain divisions; telencephalon (orange), diencephalon (purple), mesencephalon (light blue) and rhombencephalon (light green). (**b)** Volumetric comparisons of the dorsal diencephalon (green) and hypothalamus (orange). (**c)** Volumetric comparisons of the habenula, ventral thalamus and dorsal thalamus of the dorsal diencephalon, and preoptic, rostral zone, intermediate zone, and caudal zone, of the hypothalamus.(**d**) Colorimetric model depicting size differences in brain regions between surface fish and cavefish. A larger volume in surface fish results in blue coloration, while a larger volume in cavefish results in a red coloration. Horizontal optical sections depicting (**i**) dorsal, (**ii**) medial and (**III**) ventral views of the brain. Scale bars = 80 μm (a), 25 μm (b).

### Analysis of hybrid animals defines neuroanatomical associations brain-wide

Hybridization between surface and cave populations provides a powerful model for determining how high genetic diversity of natural populations contributes to phenotypic diversity. To define the volumetric anatomical relationship between wildtype populations and larval offspring, we quantified regional volume for each brain region of surface, cave, and surface to cavefish F_1_ and F_2_ hybrids. Hybrid brain regions show variability that appears consistent with different modes of inheritance, including surface dominant, cavefish dominant and surface to cavefish intermediate anatomical forms (Supplementary Data Figure 4, Supplementary Tables 41-52). We then investigated regional variability in surface to cave F_2_ hybrid brain regions by further segmenting down to local sub-nuclei (Supplementary Data Figure 5, Supplementary Tables 53-68; Supplementary Data Figure 6, Supplementary Tables 69-88). These analyses revealed molecularly defined regions that account for segment variability, such as the ventral sub-nuclei of the pallium and optic tectum. These sub-nuclei now provide discrete targets that can be functionally interrogated in future studies.

### Covariation of F2 hybrid brain regions reveals a brain following both concerted and mosaic evolution

To determine which brain regions covary, we looked for pairwise anatomical relationships across all sub regions of the brain. These results were then run through a cluster analysis to gain insight into brain-wide evolutionary mechanisms driving anatomical change in cavefish brains. Clusters of brain regions represent positive (regions are phenotypically smaller or larger together) or negative (e.g. one region is larger while the other is smaller) relationships between neuroanatomical variation (Figure 3a). This clustering analysis revealed six large clusters of neuroanatomical regions (Figure 3b), with each cluster showing strong positive anatomical relationships among subregions in that cluster. Surprisingly, we also found strong negative correlations between cluster groups (Figure 3b), suggesting that these regions have the potential to co-evolve by similar genetic mechanisms, with one group getting larger as the other gets smaller. This first analysis suggests that small sub-regions of the brain are clustering as larger modules with shared positive volumetric relationships. Alternatively, these larger modules are negatively associated with each other, suggesting a concerted evolutionary model, with large scale developmental constraints on neuroanatomical change.

**Figure 3.**
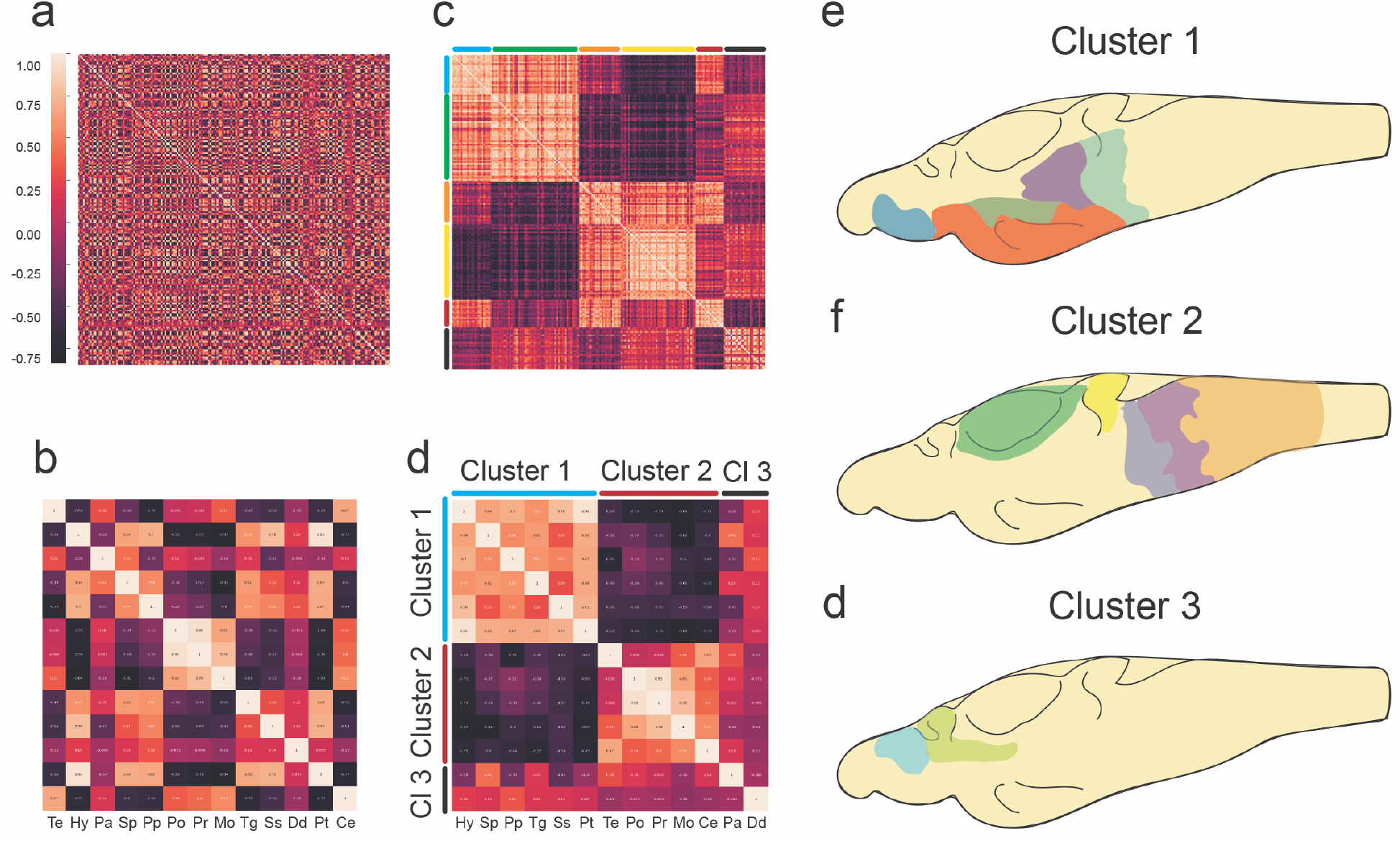
Covariation of brain region size reveals developmental tradeoff between relationships dorsal-ventral clusters brain-wide. Pairwise correlation matrix comparing covariation between (**a)** all 180 brain segments and (**b**) 13 developmentally defined regions. Clustering of pairwise correlation matrices produced **c** 6 clusters of brain segments and **d** 3 clusters of brain regions that share covariation relationships. (**c**) is the clustered matrix from (**a**), while (**d**) is the clustered matrix from (**b**). Illustrations depicting clustered segments: (**e**) cluster 1 includes the subpallium (blue), hypothalamus (orange), posterior tuberculum (olive green), tegmentum (orchid purple) and prepontine (light blue) (**f**) cluster 2 includes the optic tectum (green), cerebellum (yellow), pons (light purple), reticulopontine (orchid purple) and medulla oblongata (light orange) (**g**) cluster 3 includes subpallium (sky blue) and dorsal diencephalon (yellow green).

To help map these brain-wide relationships to larger developmental regions, we reduced our segmentation to 13 ontologically defined regions (e.g., hypothalamus, cerebellum, etc.), we then performed pairwise correlation and cluster analysis on our developmental segments for F_2_ hybrids (Figure 3c&d). This developmental cluster analysis revealed three clusters (Figure 3d), with positive associations between neuroanatomical areas within a cluster. Moreover, the volumes of regions in the two largest clusters were negatively correlated with one another, suggesting that there may be a tradeoff in the evolution of the *A. mexicanus* brain, where some areas become reduced in size at the expense of other areas increasing volumes. We then mapped neuroanatomical regions with the clusters back on the brain and found that loci within each cluster were physically localized together. Cluster one was comprised of the dorsal and caudal areas of the brain (e.g. optic tectum and medulla oblongata, Figure 3e), cluster two was predominantly made up of the ventral brain (e.g. hypothalamus and subpallium, Figure 3f), and cluster three comprised the forebrain and telencephalon (Figure 3g). These results utilized hybrid genetic variation to identify a large-scale trade-off in anatomical evolution, suggesting that early patterning events are likely modified in cavefish that result in dorsal contraction and ventral expansion.

### Geometric morphometrics provide an analytical tool for understanding the relationship between shape and volume during brain-wide evolution

Previous studies examining variation in the brain have mostly focused on volume or shape, with few providing a comparison of how shape and volume vary brain-wide (1, 2, 6, 14, 31). We sought to examine whether shape variation follows similar patterns as volume, suggesting shared origins of variation, or whether shape and volume were unrelated. To determine morphological variation in shape across the brain, we employed shape-analysis approaches previously used in assessing morphometrics of whole brain and brain regions (28, 32, 33). We first examined whether shape showed variation between populations for regions with no variation in volume, then how volume and shape relate within specific regions and finally whether shape variations follow the same brain-wide patterns seen in volumetric variation.

To begin evaluating how shape varies between surface fish and cavefish brains, we chose to characterize the pineal and preoptic region because they show no volumetric variation across populations yet play functional roles in behaviors that are highly variable across *Astyanax* populations. We found significant differences in shape of the pineal and preoptic region across wildtype surface fish and cavefish (Supplementary Table S89&90). Pachón cavefish populations exhibited a shallow preoptic region, with a wide and long pineal (Supplementary Figure S7&8). In contrast, surface fish exhibited a thin and long preoptic region, with a short and deep pineal (Supplemental Figure S7 & S8). Similar to our hybrid volumetric analysis, we then assessed shape of preoptic and pineal in surface to cave F_2_ hybrid larvae to determine how shape variation relates directly to parental populations. Shape variation of these areas in hybrids is characterized by a wide and short preoptic region, with a long, thin, and shallow pineal (Supplementary Figure S7 & S8). Importantly, surface to cave hybrids exhibit phenotypes that suggest genetically dominant (pineal, Supplementary Figure S8) and additive (preoptic, Supplementary Figure S7) modes of inheritance, suggesting that the differences in shape may be driven by genetics. This analysis suggests that regional brain shape is changed across evolution, and that these functionally important brain regions do show anatomical variation that likely impacts adaptive behaviors discovered in previous studies.

### Shape and volume variation follow the same covariation pattern brain-wide suggesting shared developmental mechanisms of brain evolution

We sought to compare brain shape variation across the brain to better understand the functional relationship between shape and volume in an evolving brain. To determine whether shape and volume was modulated by distinct or similar mechanisms, we chose to analyze two regions from our volumetric cluster 1, the optic tectum and cerebellum and two regions from cluster 2, the hypothalamus and tegmentum (Figure 4a&b). These regions allowed us to test whether shape exhibits covariation patterns similar to volume, including positive relationships within and negative relationships across dorsal and ventral clusters. First, we ran a principal component test to determine shape variation and what features were driving shape variation across hybrid individuals (Supplemental Data, Supplemental Table 92). Next, we reduced PC1 to a single value, to provide a pairwise correlation and cluster analysis. This process allows us to gauge how shape variability relates across brain regions, testing whether variation in shape and volume are modified by the distinct or varying developmental mechanisms.

**Figure 4.**
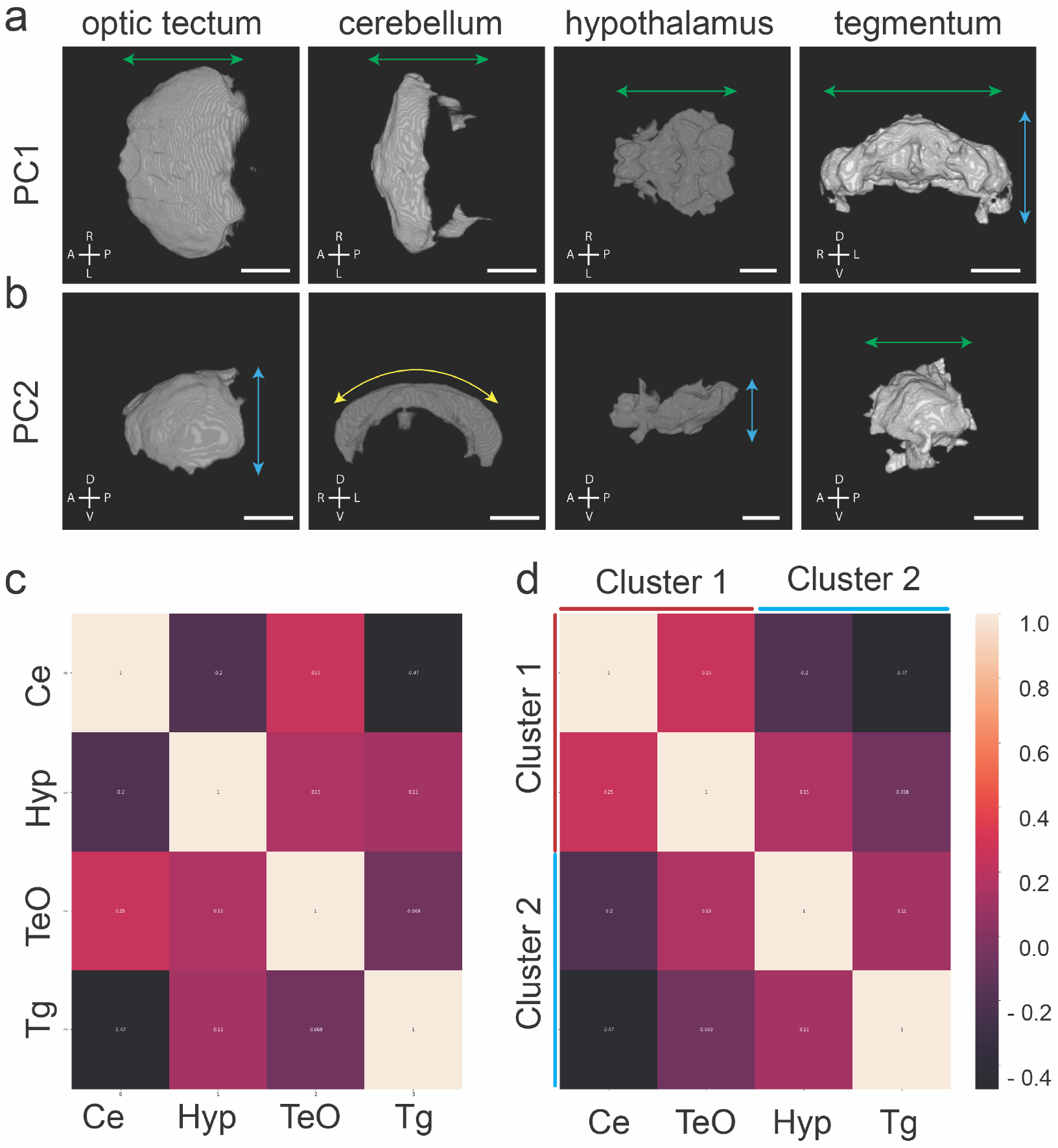
Shape covariation suggests volume and shape share brain-wide mechanism of brain evolution. Representatives of shape analysis and vector arrows for (a) principal component 1 (PC1) and (b) principal component 2 (PC2). Arrow colors denote dimension; green = length, blue = depth, yellow = degree of curvature. (c) Correlation matrix comparing the covariation relationship between regions from volumetric covarying cluster 1, cerebellum (Ce) and optic tectum (TeO), and cluster 2, hypothalamus (Hyp) and tegmentum (Tg). (d) A cluster analysis of covariation grouped regions into 2 clusters as predicted by volumetric covariation. Scale bars = 100 μm.

Finally, to test whether anatomical variation in shape and volume across the brain follow similar dorsal-ventral trade-offs, we applied our PC1 values from our shape PCA and performed a correlation and cluster analysis to determine if anatomical shape covaries the same as volume for hybrid brain regions. We found that covariation of anatomical shape clusters the same as volume in a dorsal-ventral fashion, with cluster 1 and cluster 2 showing positive relationships within clusters, and negative relationships across clusters (Figure 4c&d). Our hybrid shape analysis shows that mechanisms of concerted brain evolution are impacting both anatomical volume and shape to reorganize the dorsal-ventral development of cavefish brains.

## Discussion

Here we establish a laboratory model of anatomical brain evolution that utilizes an innovative molecularly defined neuroanatomical atlas and applied computational tools, which can be used to assess how all brain regions have evolved anatomically. The ability to apply this atlas and the computational approaches to F_2_ hybrid fish permits a brain-wide dissection of how neuroanatomy changes and a powerful analysis of not only how different neuroanatomical areas evolve but also which areas co-segregate together. These studies reveal that the *Astyanax* brain has evolved through both evolutionary models of anatomical evolution, via developmental constraint with a brain-wide dorsal-ventral relationship, while also experiencing a local functional constraint between the pallium and thalamus, regions with well-established functional dependence for development and maturation (21, 35, 36). Finally, this study is one of the first to directly assess how the volume and shape of regions relate to one another across all regions of the brain.

Previous studies examining how the brain evolves have largely been restricted to comparative studies between closely related, albeit different species, and these studies have revealed gross differences in neuroanatomy, connectivity, and function between derived animals (5, 9, 10, 12, 14). However, the *Astyanax* model provides a powerful tool for assessing how the brain evolves in a single species with multiple divergent forms and an extant ancestor (20, 22, 37). Moreover, because surface and cave forms are the same species, the ability to produce surface/cave and cave/cave F_2_ hybrid fish permits a powerful dissection of functional principles underlying brain evolution (25, 26). We previously published population specific neuroanatomical atlases for this species and used this to comparatively examine how gross neuroanatomy and physiology relate to behavior in pure surface and cavefish (29, 30). The current study adds several powerful principles to the ongoing work utilizing this model to understand how the brain evolves, including a single atlas for all populations to functionally compare neuroanatomy in pure and hybrid offspring, automated brain segmentation for 180 annotated sub-populations of neurons, and the application of computational approaches for a complete whole-brain assessment of the evolution of the brain.

Two prominent theories suggest that either the majority of the brain is impacted by allometry (scaling with body size) and the constraint of shared developmental programs, and the mosaic hypothesis, which says that more discrete regions will independently evolve based on shared function (7, 8). Our data provides evidence that both evolutionary mechanisms are utilized to drive anatomical evolution of cavefish brains. First, a concomitant relationship between dorsal-caudal shrinkage and rostral-ventral expansion exemplifies concerted evolution, while independent expansion of the pallium represents mosaic evolution. Our data show that the dorsal-caudal areas of the brain evolve together, and that regions such as the optic tectum and the cerebellum, two areas that constitute a large proportion of the dorsal-caudal region shrink in size. In contrast, rostral-ventral areas coevolve together, such as the hypothalamus and subpallium that are enlarged in cavefish. Importantly, we find in F_2_ hybrid fish that reduced optic tectum and cerebellum are concomitate with an enlarged hypothalamus and subpallium, suggesting that expansion of some regions come at the expense of others. Whether these anatomical relationships exist in other independently evolved cavefish populations is currently unknown, and future studies in these populations would provide more robust support for evolutionary patterns.

In addition to volume, evolutionary changes in shape of neuroanatomical regions have been shown to alter function of different regions. The mammalian cortex, for example, has evolved from a smoother lissencephalic cortex in more ancestral species to a folded one in higher order animals such as primates (7, 38, 39). Folding of the cortex is thought to increase surface area and has been implicated in more complex processing of the brain (40, 41). Whether such changes in shape are important for non-cortical areas, or the relationship between shape and volume is not known. However, we do know that shape variation has been shown to be a common adaptation in other tissues. Beak differences in Galapagos finches have been shown to change in accordance with the size food sources, and such changes have been shown to rely on differences in bone morphogenic protein signaling (42, 43). Craniofacial differences in African cichlids also have been shown to vary as an adaptive quality to food availability (32, 33). Furthermore, standard methods for assessing complex shape features have been applied to studying brain shape evolution in non-model organisms, generating anatomical evolutionary hypotheses that have lacked an appropriate model for assessing functional mechanisms of anatomical evolution (9, 10, 34). By applying these morphological measuring and analyzing methods with our hybrid volume pipeline, we were able to see that complex shape phenotypes are likely genetically encoded, evidenced in hybrid intermediate phenotypes, and that shape and volume variation in developmentally constrained clusters are impacted by shared mechanisms. While, the functional and adaptive significance of differences in shape are not known, future work relating neuronal activity and function with differences in shape in this model could help address this question.

Together, these studiessupport general principles underlying the evolution of the vertebrate brain. This study represents the first computational brain atlas for a single species with multiple evolutionary derived forms, and the application of the atlas to hybrid animals represents the first assessment of how different neuroanatomical areas evolved in both volume and shape. Moreover, by combining this atlas with myriad cutting edge tools that we have generated for this model, functional neuroimaging, and genome editing tools, will allow researchers to identify the genetic mechanisms that explain these changes. The strong genetic and neuronal conservation of the vertebrate brain, as well as the simplified nervous system of fish, suggests that this model offers great potential to understand general principles of neuroevolution.

## Materials and Methods

### Fish maintenance and husbandry

Mexican tetras (*A. mexicanus*) were housed in the Florida Atlantic Universities Mexican tetra core facilities. Larval fish were maintained at 23**°**C in system-water and exposed to a 14:10 hour light:dark cycle. Mexican tetras were cared for in accordance with NIH guidelines and all experiments were approved by the Florida Atlantic University Institutional Care and Use Committee protocol #A1929. *A. mexicanus* surface fish lines used for this study; Pachón cavefish stocks were initially derived from Richard Borowsky (NYU); Surface fish stocks were acquired from Texas stocks. Surface Rio Choy were outcrossed to Pachón to generate F_1_ hybrids, while F_1_ hybrid offspring were backcrossed to produce F_2_ hybrids.

### Immunohistochemistry and imaging

Larval immunohistochemistry was performed as previously published [Kozol et al. 2021], using antibodies raised against total ERK (ERK; p44/42 MAPK (ERK1/2), #4696) and Islet-1 & Islet-2 homeobox (Islet1/2, #39.4D5, Developmental Studies Hybridoma Bank, University of Iowa, Iowa City, IA; Supplemental Table S71). IHC stained larvae were imaged on a Nikon A1R multiphoton microscope, using a water immersion 25x, N.A. 1.1 objective.

### Automated segmentation and brain region measurements

To segment and measure subregions of the brain, we modified the zebrafish brain browser brain atlas, neuropil and cell body mask for the existing zebrafish resource CobraZ by using previously published Advanced Normalization Toolbox (ANTs; (27, 44)) registration and inverse registration scripts. A surface to Pachón F_2_ hybrid larval brain was stained with ERK, registered and inverse registered (automated segmentation) to the zbb ERK standard brain, resulting in our hybrid ERK standard brain. This produced the *Astyanax* brain atlas that was then used for automated segmentation of all ERK stained brains for all *Astyanax* populations. CobraZ measures the size of segmented regions of the brain and calculates regional size as percent of total brain (pixels of brain region/total pixels in brain (27)). To test the accuracy of our modified version of CobraZ, we hand segmenting 20 brains each for Surface fish, Pachón cavefish and F_2_ surface to cave hybrid larvae. These labeled neuroanatomical areas were then compared to automated segmentation from our brain atlas by running a custom cross correlation script. 3D volumetric images were imported into matlab using the ‘imread’ function, vectorized to a 1D vector using ‘imreshape’, and then a Pearson’s correlation was performed using the ‘corr’ function (scripts available upon request). This cross-correlation analysis revealed >80% correlation between ERK defined hand and automated segmentation, with no significant differences in the variation of correlation across populations (Supplementary Figure S2b, Supplementary Table. S1). In addition, we produced a modified segmentation file that defines larger subregions that overlap with tERK neuropil to provide cross correlation analysis across brain regions and populations. Finally, we tested the accuracy of the cerebellum and hypothalamic subregions, using the Islet1/2 antibody that labels previously described discrete populations of cell bodies (45)(Supplemental Material, Table 1).

### Pairwise correlation and covariation analysis of brain region volumes and shapes

Correlations between volumes of brain regions were determined using custom written scripts in python. Volume data was imported from Microsoft Excel into Python using the pandas library. Scipy was then used to determine the pairwise correlation between all brain regions. The seaborn library was then used to generate a heat map with annotations set to “True” to overlay correlation coefficients on the pairwise correlation matrix. Cluster analysis of the corresponding pairwise correlation matrix was performed using scipy toolkit. The distance matrix was first calculated from the correlation matrix and then indexed into the corresponding clusters. The correlation matrix was then clustered by grouping all regions that clustered (i.e., had the same index value). The resulting metric was again generated using seaborn. Detailed Jupyter notebooks will be made available upon request from the corresponding authors.

### Brain segment geometric morphometric analysis

After extracting 3D models of the preoptic region of the hypothalamus and pineal body of the diencephalon, we characterized shape variation in the Preoptic Region and Pineal body of Pachón (n=24), surface (n=16), and F_2_ hybrid populations (n=34) using 3D geometric morphometrics (LandmarkEditor (v3.0) (46). To assess differences in the preoptic region, we placed 16 landmarks across the preoptic region, performed a procrustes superimposition on our shape data to remove the effects of translation, rotation, and scaling from all individuals using the gpagen function from the geomorph (v4.0) package in R (47, 48). We then assessed differences in shape among populations and performed a multivariate regression of shape on centroid size and population (i.e., surface, cave, hybrid) using the procD.lm r function from geomorph. Unlike other sub regions of the brain, the pineal body represented a shape with few obvious homologous points to place landmarks. For the pineal body, we placed two fixed landmarks at the anterior and dorsal apexes of the pineal and surrounded the base of the pineal with 26 sliding semi-landmarks. We then took advantage of a procedure to automate the placement of 99 surface landmarks across the pineal region to wrap the pineal body with sliding surface semi-landmarks to best characterize the shape of this sub region among individuals. This required building a computer aided design (CAD) template of the pineal using FreeCAD (v.0.16.6712), which we modeled as a hemisphere, and placing the fixed landmarks, sliding semi-landmarks (Supplemental Figure. S8), and surface landmarks on the CAD model using LandmarkEditor. We then used the *R* package Morpho to map the surface landmarks from the template to the pineal model of each individual specimen using the placePatch function (49, 50). We also performed a principal component (PC) analysis on our shape data to visually examine the major axes of shape differentiation among populations.

To assess the degree of association between brain sub region shape and volume we performed a multivariate regression of shape on volume using the procD.lm r function from geomorph. Similarly, to assess associations among brain sub region shapes we performed a partial least squares (PLS) analysis using the two.b.pls r function from geomorph.

### Statistics

All wildtype population standard t-tests were calculated using the program CobraZ. For hybrid population comparisons, Prism (Graphpad Software Inc., San Diego, CA) was used to run standard ANOVAs, followed by a Holm-Šídák’s multiple comparisons test. To evaluate covariation of F_2_ subregions, geometric morphometry analyses were all conducted in R (47, 48, 50) using the packages geomorph (v4.0) and Morpho (v2.6) (47-49) to assess associations and produce morphospace plots.

## Supporting information

Supplemental Data Full

## Data Sharing

All raw and analyzed data, custom code and adapted tools will be made available upon request from the corresponding authors.

## Acknowledgements

We would like to thank Dr. Harry Burgess for his help in adapting the zebrafish brain browser atlas and modifying files for CobraZ to analyze *Astyanax mexicanus* neuroanatomy. We thank Dr. James Jaggard for his expertise and early work on the formation of this project. To the administration and staff at the Jupiter Life Science Initiative in the Department of Biology at Florida Atlantic University, especially Peter Lewis and Arthur Lapatto for overseeing the health and care of the FAU *Astyanax* fish facility.

## Supplementary Material

### Supplementary Data

#### A single atlas for studying differences in neuroanatomy across surface fish, cavefish and hybrid populations

To characterize brain wide neuroanatomical difference between *A. mexicanus* populations, isolated surface *Astyanax* from the Rio Choy population as well as cavefish from the Pachón cave population. Cavefish have several phenotypes that distinguish them from surface fish, including loss of pigmentation and eyes. These phenotypes can also be seen in larval fish. The small size of the larval fish permits brain wide imaging, and thus all experiments were done in 6 dpf larvae (Fig 1a). To produce a segmented atlas for cave and surface Astyanax, we used a previously published neuroanatomical atlas from a related fish, the common zebrafish (*Danio rerio*)^1,2^, that is neuroanatomically homologous with *A. mexicanus*^3^. The zebrafish brain browser (zbb) brain atlas was constructed using 210 unique neuroanatomical markers, that culminated in the segmentation of 180 different brain regions^4-6^. To adapt the zbb atlas, we stained brains from four groups of *A mexicanus* fish: surface fish, Pachòn cavefish, Surface/Pachón F_1_ and F_2_ hybrids. Brains were stained with total ERK antibody, a brain-wide neuroanatomical marker validated in both zebrafish and *A. mexicanus*^3,7^ (Supplemental Data, Figure S3b). Next, we used an image normalization program, advanced normalization toolbox^8,9^ (ANTs), to overlay or register our standard surface to cavefish F_2_ hybrid brain to the standard zbb atlas brain (Supplementary Data Figure S3c). This process provided a set of instructions that could be reversed to map the zbb segmented atlas onto our hybrid standard brain (Figure 1b). This created a hybrid brain atlas that could be used to register brains from all four *A. mexicanus* populations, producing a single computational atlas for measuring brain size and shape (Supplemental Figure, S3c&d). We validated our *Astyanax* segmented atlas using two distinct approaches. First, we assessed registration efficiency using a cross-correlation analysis between registered *Astyanax* brain and the zbb reference brain. These data revealed that the two were highly correlated (rho=0.95, Supplemental Tables), suggesting a high level of overlap among pixels and highly efficient registration procedure. In order to confirm that segmentations from zbb accurately predicted neuroanatomical areas in *Astyanax*, we labeled a subset of brains with an antibody that labels distinct neuroanatomical loci (Fig. 1d). The Insulin gene enhancer protein ISL-1 and Insulin gene enhancer protein ISL-2 (Islet1/2) antibody labels clusters of neurons across the brain, including the preoptic region, and rostral, intermediate and caudal hypothalamic regions. Knowing that Isl1/2 neurons are located in these brain segments, we predicted that reverse registration of our F_2_ cavefish atlas onto larvae would automatically segment these neuronal clusters into the correct brain region. As predicted, registered *Astyanax* brains labeled with islet1/2 had fluorescence staining that co-localized in segments of the subpallium, hypothalamus and hindbrain, suggesting that the zbb segmentation accurately labeled the *Astyanax* brain atlas (Figure. 1d, Supplementary Figure S3). Together, these data provide a well annotated brain atlas for *Astyanax mexicanus* that permits brain-wide comparisons of regional anatomy across anatomically distinct populations of a single species.

#### Automated volumetric comparisons of brain regions reveal highly localized changes in functionally defined brain segments

Our atlas segments 180 neuroanatomical regions that can be combined to scale from major subdivisions of the brain to more functionally relevant units. To look at the major subunits, we combined regions into the telencephalon, diencephalon, mesencephalon and rhombencephalon. While overall brain size was larger in cavefish than surface fish, the major brain divisions and their segments vary in comparative size (Supplementary Figure S3). Consistent with manually segmented brains, the mesencephalon was significantly smaller in cavefish compared to surface fish. By contrast, the telencephalon, diencephalon and rhombencephalon were all larger in cavefish relative to surface animals (Fig. 2a, Supplementary Figure S4).

#### Hybrid anatomical variation can be used to target discrete subnuclei for functional studies

To define volumetric variation in molecularly segmented regions of interest, we examined the optic tectum, a region that is known to be developmentally and anatomically divergent in cavefish and surface fish. Our results support previous work showing the optic tectum is smaller in cavefish ^10,11^. The average optic tectum size in F_2_ hybrids is also reduced relative to surface fish, with the optic tectum of most hybrid individuals similar in size to that found in cavefish (Figure 2b). However, further segmenting shows that F_2_ hybrids have highly variable volumes in the grey matter (cell bodies), specifically in the ventral bulk of the optic tectum, suggesting that phenotypic variation in molecularly segmented subregions can be masked by only analyzing larger developmentally annotated regions (Figure 2c&d). We then applied this method to determine whether we could isolate regional variation in subregions of the telencephalon, a brain region that iis vital for cognitive states from memory to emotion ^12-14^, and has been shown to be expanded in cavefish ^11,15^ (Figure 2a). We discovered that the telencephalon exhibits high phenotypic variability across all three major subregions, suggesting the most recently evolved brain region in chordates ^12^ has a rapidly expanding olfactory bulb, subpallium and pallium in cavefish (Figure 2b). A further anatomical refinement revealed that caudal segments of the olfactory bulb and subpallium, along with the ventral pallium are the main source of volumetric variability for each region, providing anatomical targets for for future functional studies (Supplementary Figure S6c, d). These results reveal that brain-wide anatomical evolution can be phenotypically resolved at a high-resolution by comparing hybrid larvae with parental populations.

#### Shape and volume relationships within regions are governed by local developmental constraints

First, we analyzed differences in shape across F_2_ individuals for each region, to provide a description of how the shape varied in different axes (Supplemental Data, Table S2). Then we assessed the degree of association between volume and shape within and across regions. We performed a principal component analysis to determine whether shape varied and what shape features were driving shape variation within the F2 hybrid population. We found that all regions showed significant shape variation, along the length of the shape for the optic tectum, cerebellum and hypothalamus exhibiting variation, and along the depth and width of the tegmentum (Figure 4a, Supplementary Table 91). We then extracted our PC1 measurements for shape and compared them to relative volumes for all four regions. We found significant associations between volume and shape in three of four brain regions (Supplementary Table 91). There were strong associations between volume and shape in the cerebellum, tectum, and tegmentum, while we found no association in the hypothalamus. These results show that within individual brain regions, variation in shape and volume can exhibit either shared or distinct mechanisms, results that likely reflect regional relationships based on developmental and physical constraints.

### Supplementary Figures

Supplementary Figure S1. Pipeline for immunohistochemistry, automated segmentation, and volumetric comparisons for individual fish larvae.

Supplementary Figure S2. Cross correlation analysis between hand and automated segmentation of total-ERK defined brain segments.

Supplementary Figure S3. Variation in segment volume between surface and cavefish populations.

Supplemental Figure S4. Volumetric variability in hybrid larvae reflect wildtype genetic diversity through dominant and intermediate phenotypes.

Supplemental Figure S5. Scalable segmentation of the tectum identifies high variability in the ventral sub-nuclei of the optic tectum’s cell layers.

Supplementary Figure S6. Hybrid brains link genetic variation in wildtype populations to anatomical variation in distinct sub-nuclei of the olfactory bulb, subpallium and pallium.

Supplemental Figure S7. Shape variability of the preoptic region in hybrid larvae display an intermediate phenotype between wildtype populations

Supplemental Figure S8. Pineal shape variation in hybrids exhibits a cavefish dominant phenotype.

**Extended Data Figure 1.**
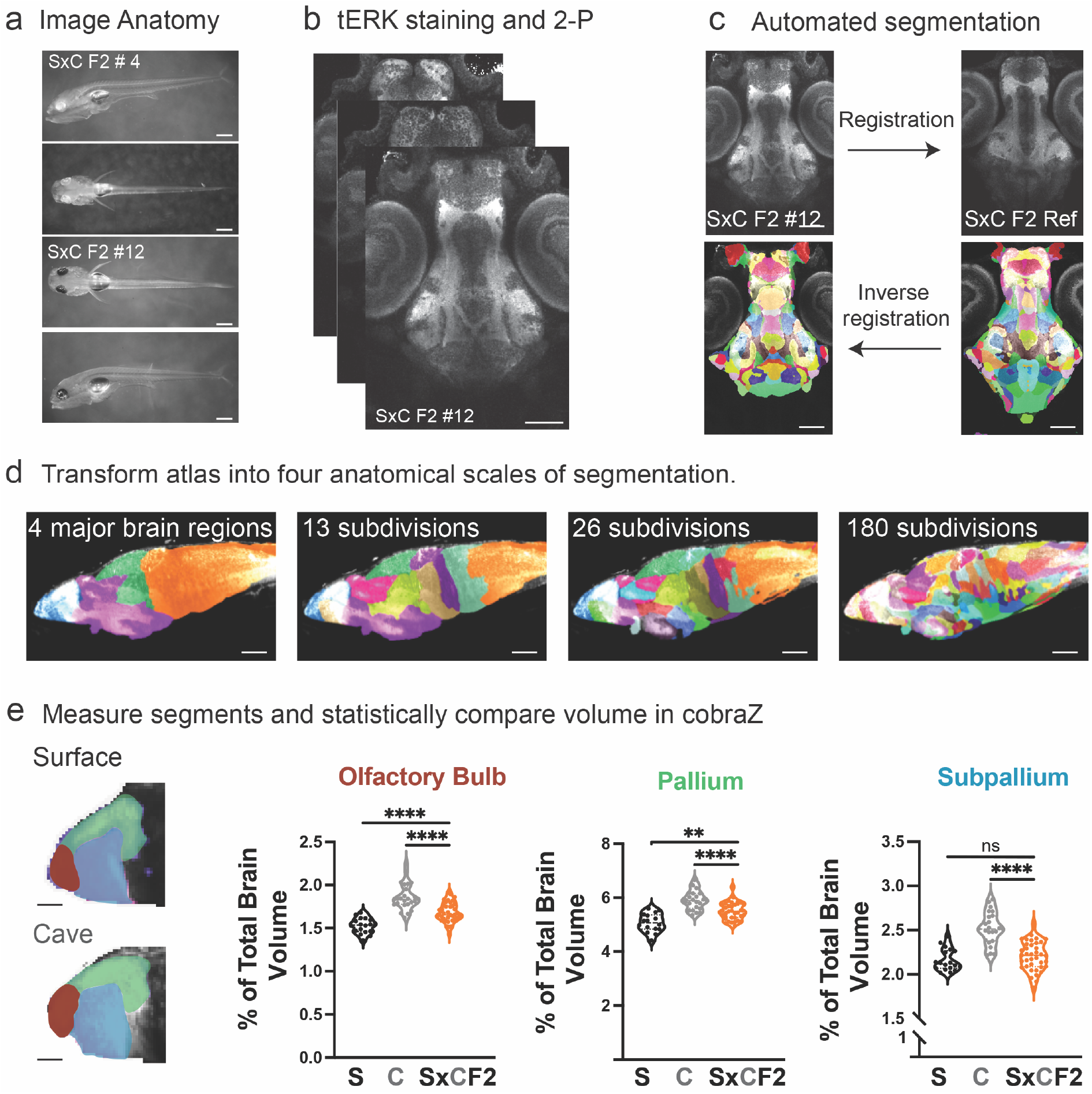
Pipeline for immunohistochemistry, automated segmentation and volumetric comparisons for individual fish larvae. (**a**) Larval fish were imaged at 5 dpf to measure external anatomical features (e.g. standard length). (**b**) Larvae were fixed at 6 dpf, immunostained with total-ERK (tERK) and imaged on a 2-photon microscope. (**c**) All larvae were registered to a surface to cave F_2_ hybrid reference brain, followed by an inverse registration of the segmented cavefish atlas. (**d**) The 180 brain segment ZBB atlas was transformed into scalable atlases, 4 major subdivisions, 13 developmentally defined and 26 molecularly and functionally defined regions. (**e**) segmented larval brains were finally run through CobraZ to volumetrically measure and statistically compare each brain region. Scale bar = a. 500 μm, b.&c. 100 μm, d. 50 μm and e. 25 μm.

**Supplementary Figure S2.**
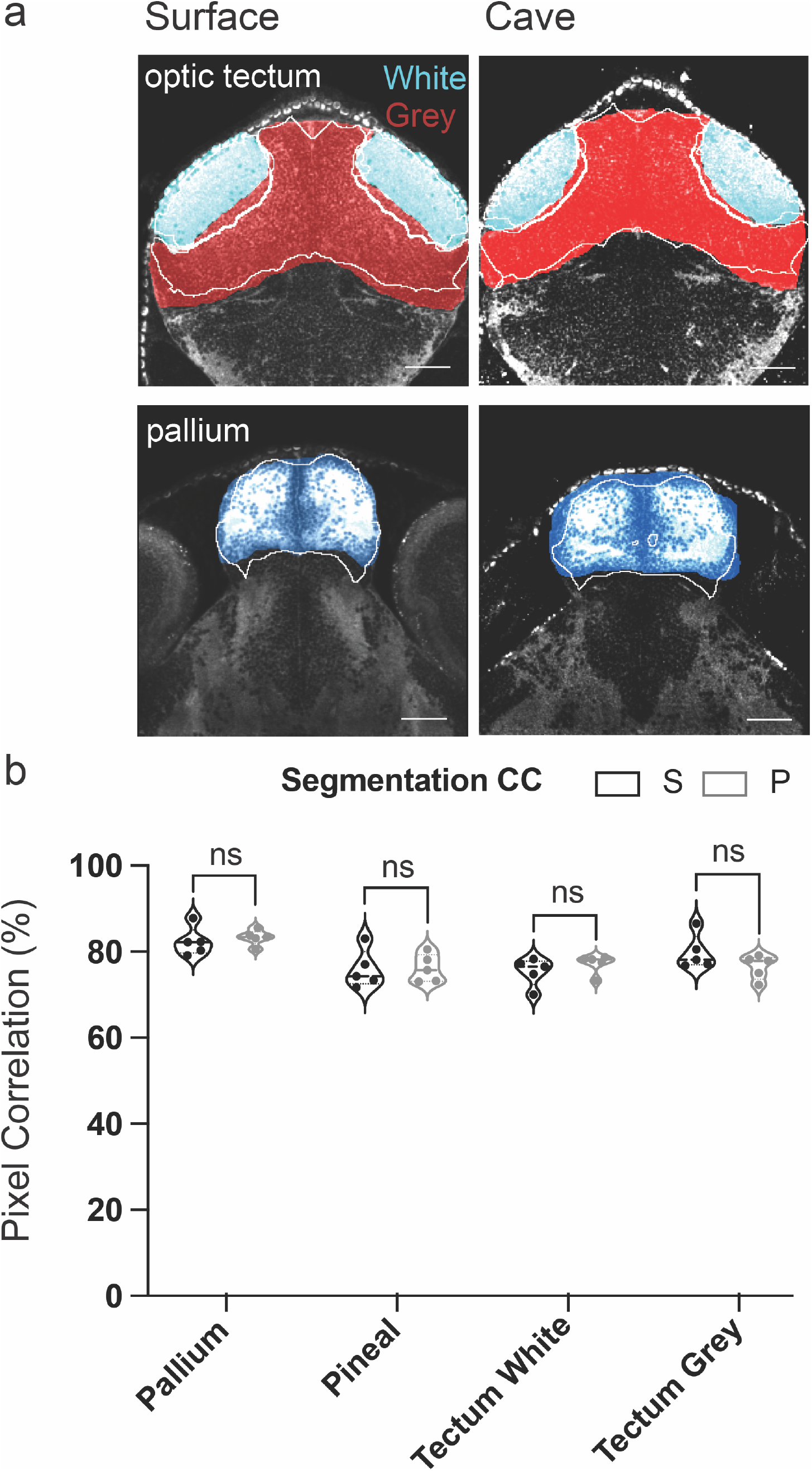
Cross correlation analysis between hand and automated segmentation of total-ERK defined brain segments. (**a**) Optical sections (coronal) through the and optic tectum (white matter light blue and grey matter red) and pallium (dark blue). Hand segmentations are in color, while automated versions are shown in white outline. (**b**) Cross correlation analysis comparing the percent of overlap between hand and automated segments for ERK defined regions. Pixel correlation percentages did not vary between surface and cavefish. Cross correlation percentages were compared using a standard t-test. Scale bars = 25 μm.

**Supplementary Figure S3.**
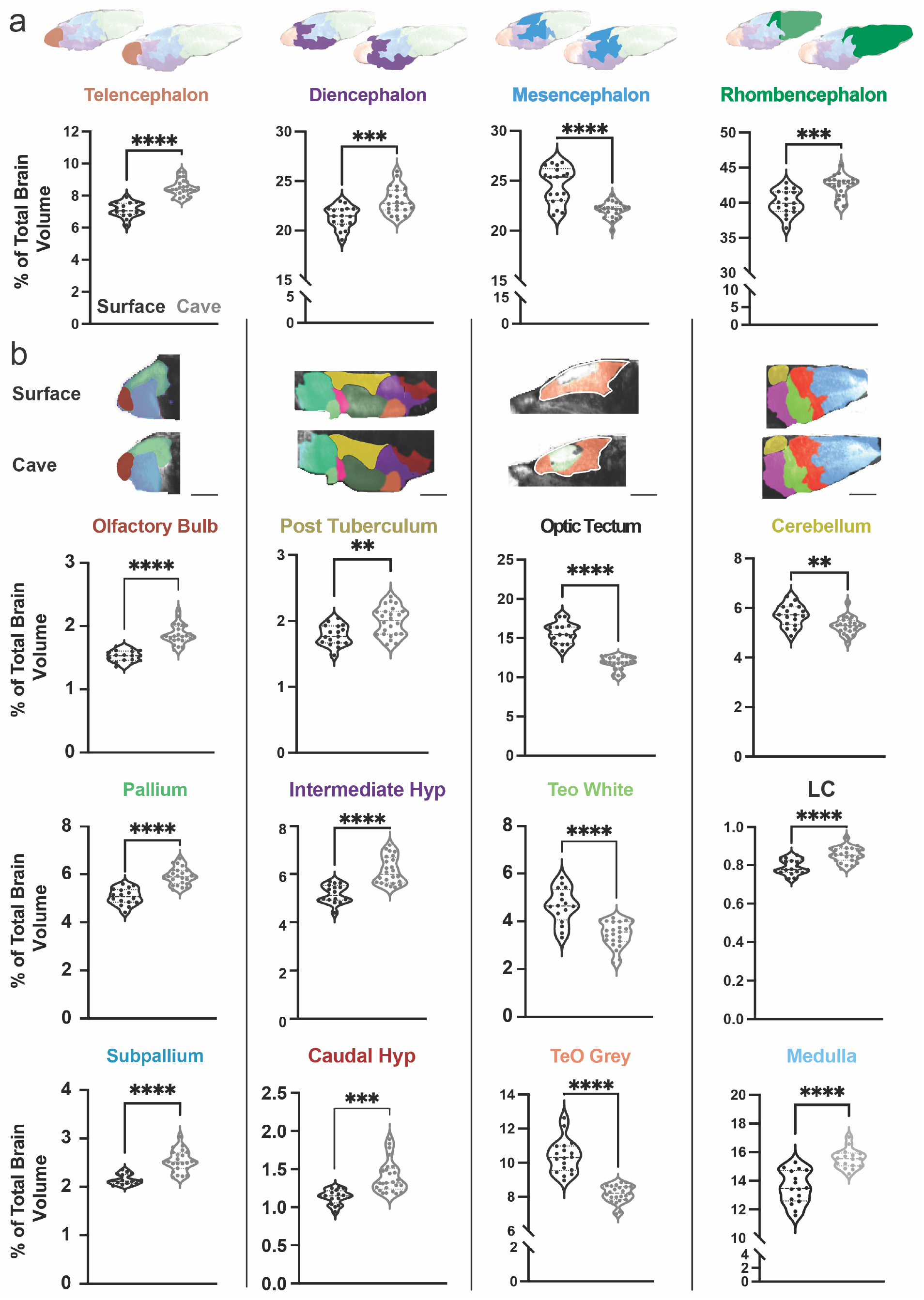
Variation in segment volume between surface and cavefish populations. (**a**) Major brain divisions of the vertebrate brain and box plots comparing the volume of each region between surface and cave. Volumes are reported as percentage of total brain volume, calculated by dividing total pixels in a segment by total pixels within the brain. (**b**) Columns segmenting each major brain division into brain regions defined by developmental, molecular and functional categories. Each box plots brain segment is color coded to the corresponding atlas picture at the top of each column. All segments were statistically analyzed using a students t test. P value significance is coded as; * = p<0.05, ** = p<0.01, *** = p<0.001, **** = p<0.0001. Scale bars = 40 μm (Tele, Dien, Mesen), 80 μm (Rhomb).

**Supplemental Figure S4.**
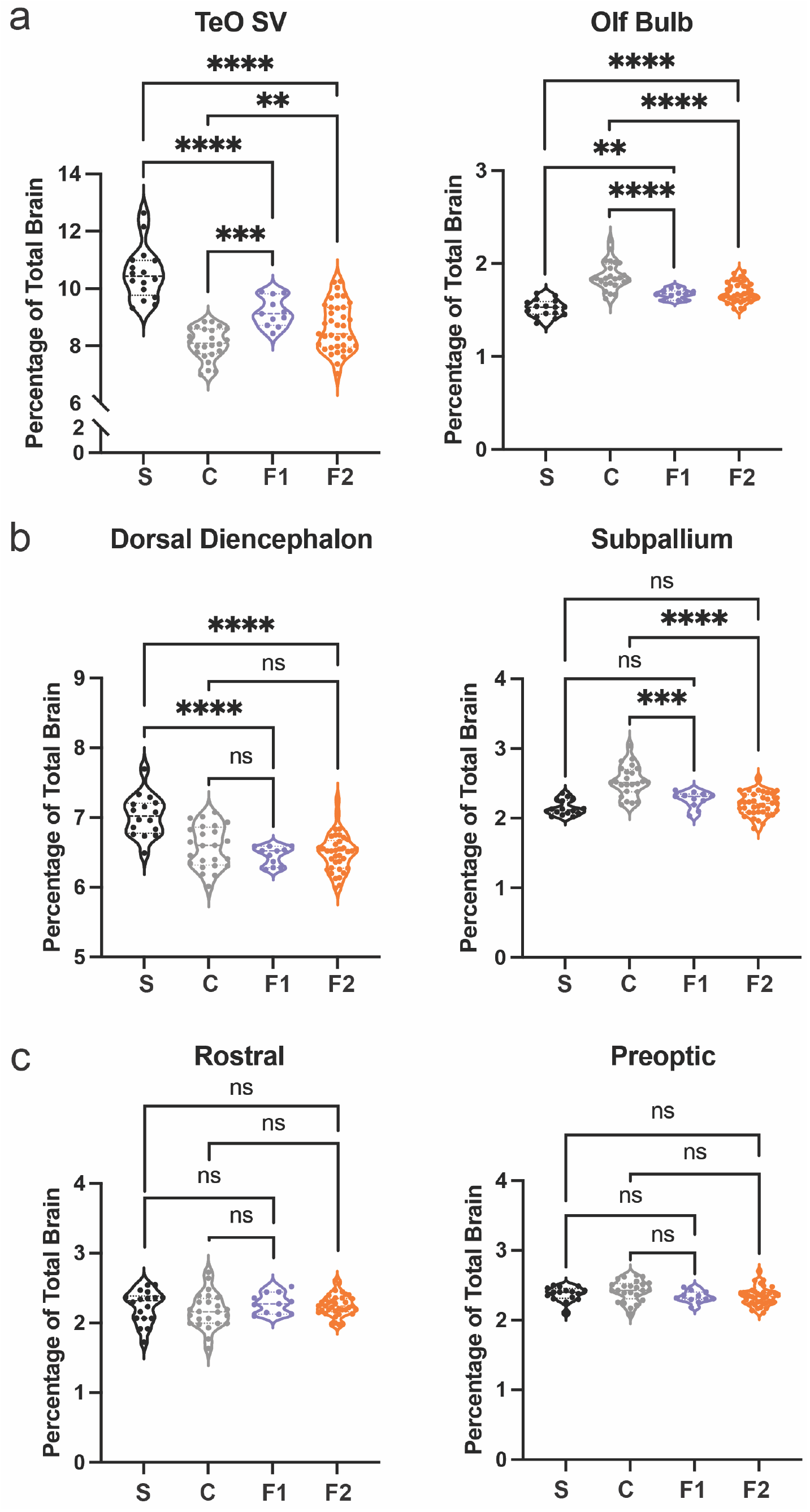
Volumetric variability in hybrid larvae reflect wildtype genetic diversity through dominant and intermediate phenotypes. Representative regions for hybrid larvae that show (**a**) an intermediate brain size between wildtype surface and cave values, (**b**) genetic dominance with Pachón cavefish (dorsal diencephalon) or surface fish (subpallium) wildtype populations, and **c** no difference between hybrid and wildtype populations.

**Supplementary Figure S5.**
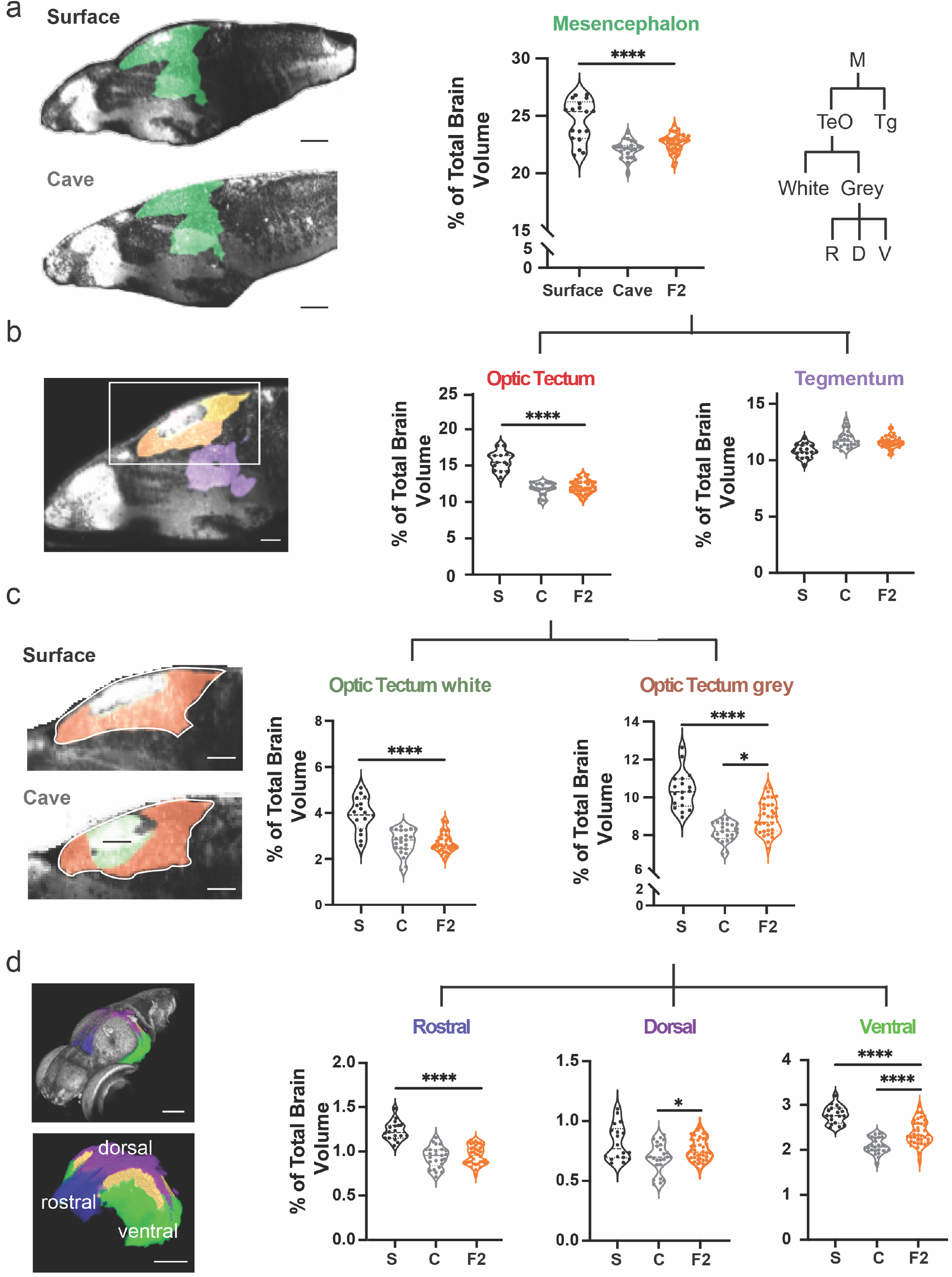
Scalable segmentation of the tectum identifies high variability in the ventral sub-nuclei of the optic tectum’s cell layers. (**a**) Volumetric comparison of the mesencephalon in surface, Pachón cavefish and surface to Pachón hybrid larvae. Sagittal sections showing the mesencephalon (green). Percent total brain volume represents pixels of segment divided by total pixels in the brain. Segment tree abbreviations, M – mesencephalon, TeO – optic tectum, Tg – tegmentum, R – rostral, D – dorsal, V – ventral (**b**) Volumetric comparisons of the optic tectum (yellow) and tegmentum (purple). (**c**) Volumetric comparisons of the optic tectum white (neuropil; forest green) and grey matter (cell bodies; orange). (**d**) Volumetric comparisons of rostral (royal blue), dorsal (purple), and ventral (lime green) segments of the optic tectum grey matter. Scale bars = 80 μm (a), 25 μm (b), 50 μm (c&d).

**Supplementary Figure S6.**
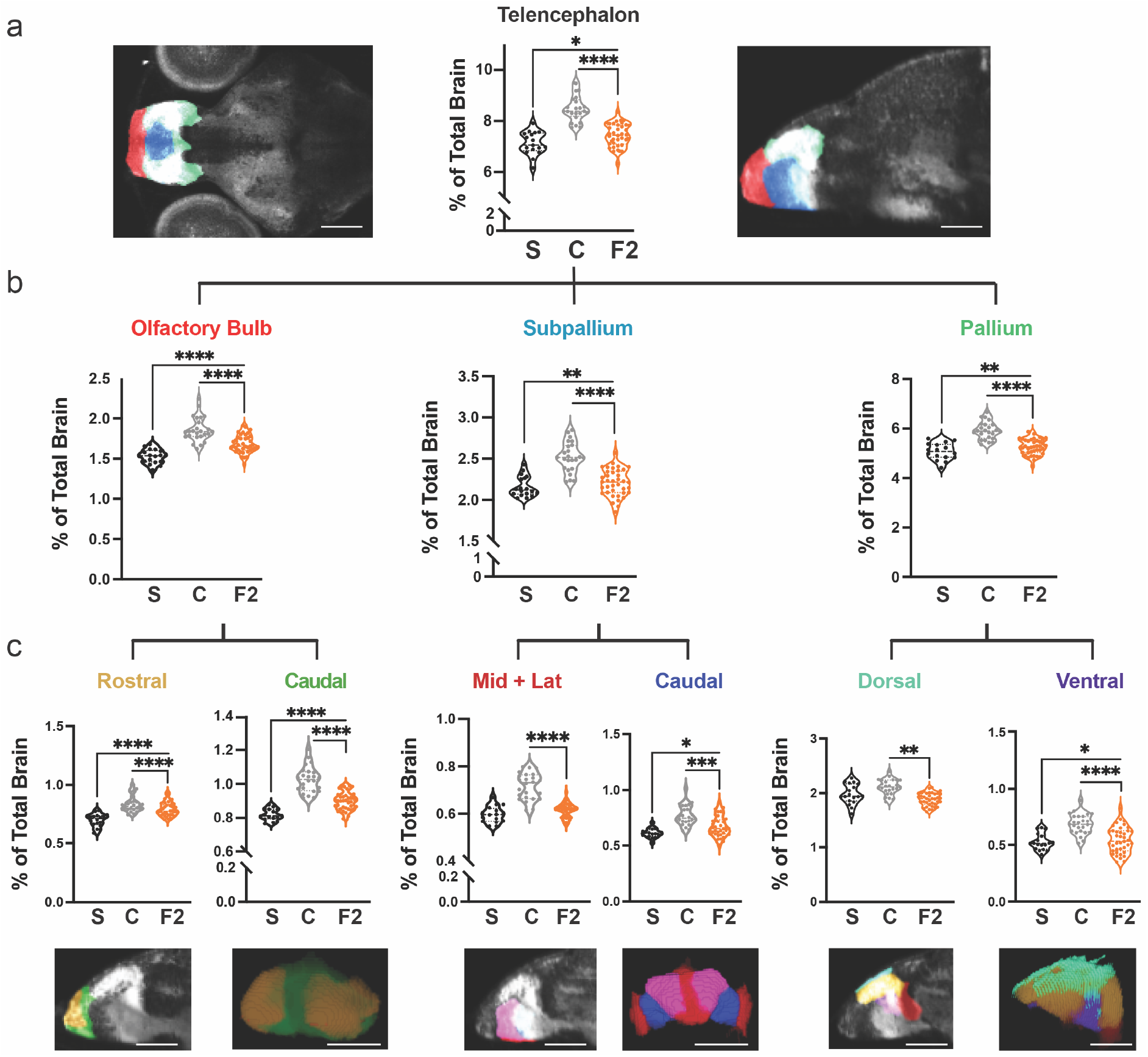
Hybrid brains link genetic variation in wildtype populations to anatomical variation in distinct sub-nuclei of the olfactory bulb, subpallium and pallium. (**a**) Volumetric comparison of the telencephalon in surface fish, Pachòn cavefish and surface to cave F_2_ hybrids. Percent total brain volume represents pixels of segment divided by total pixels in the brain. (**b**) Sagittal sections show subdivisions of the telencephalon, pallium (green), subpallium (light blue) and olfactory bulb (red). (**c**) Volumetric comparisons of discrete regions of the pallium, dorsal (sky blue) and ventral (purple), and olfactory bulb, rostral (gold) and caudal (green). Scale bar = 80 μm (a), 50 μm (c)

**Supplemental Figure S7.**
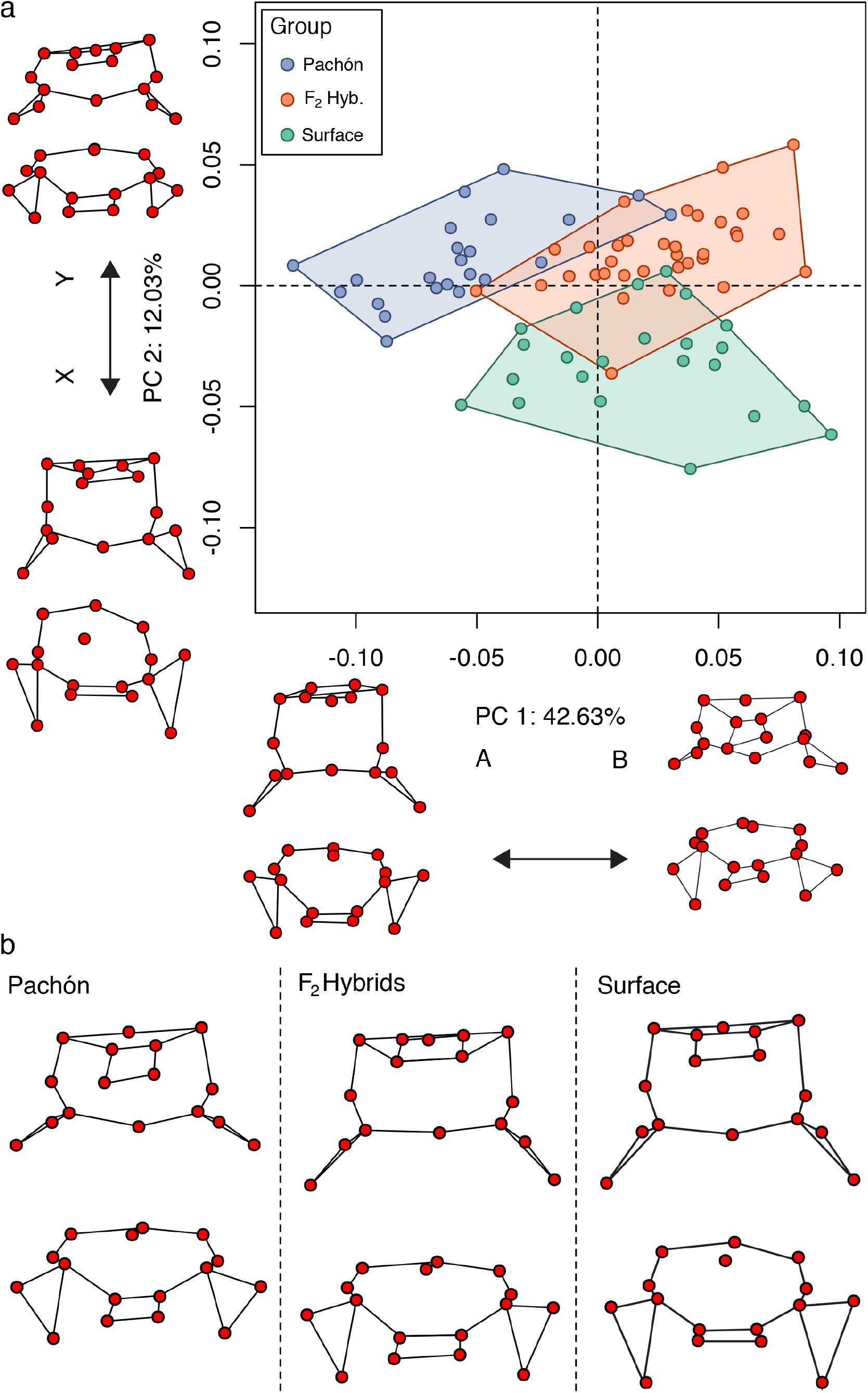
Shape variability of the preoptic region in hybrid larvae display an intermediate phenotype between wildtype populations (**a**) PCA capturing 54 percent of the variation across surface, cave, and surface to cave F_2_ hybrids. PC1 describes preoptic width, while PC2 describes length. (**b**) Illustrations of the median shape for each population. Top row provides an anterior view, bottom row provides a side view.

**Supplemental Figure S8.**
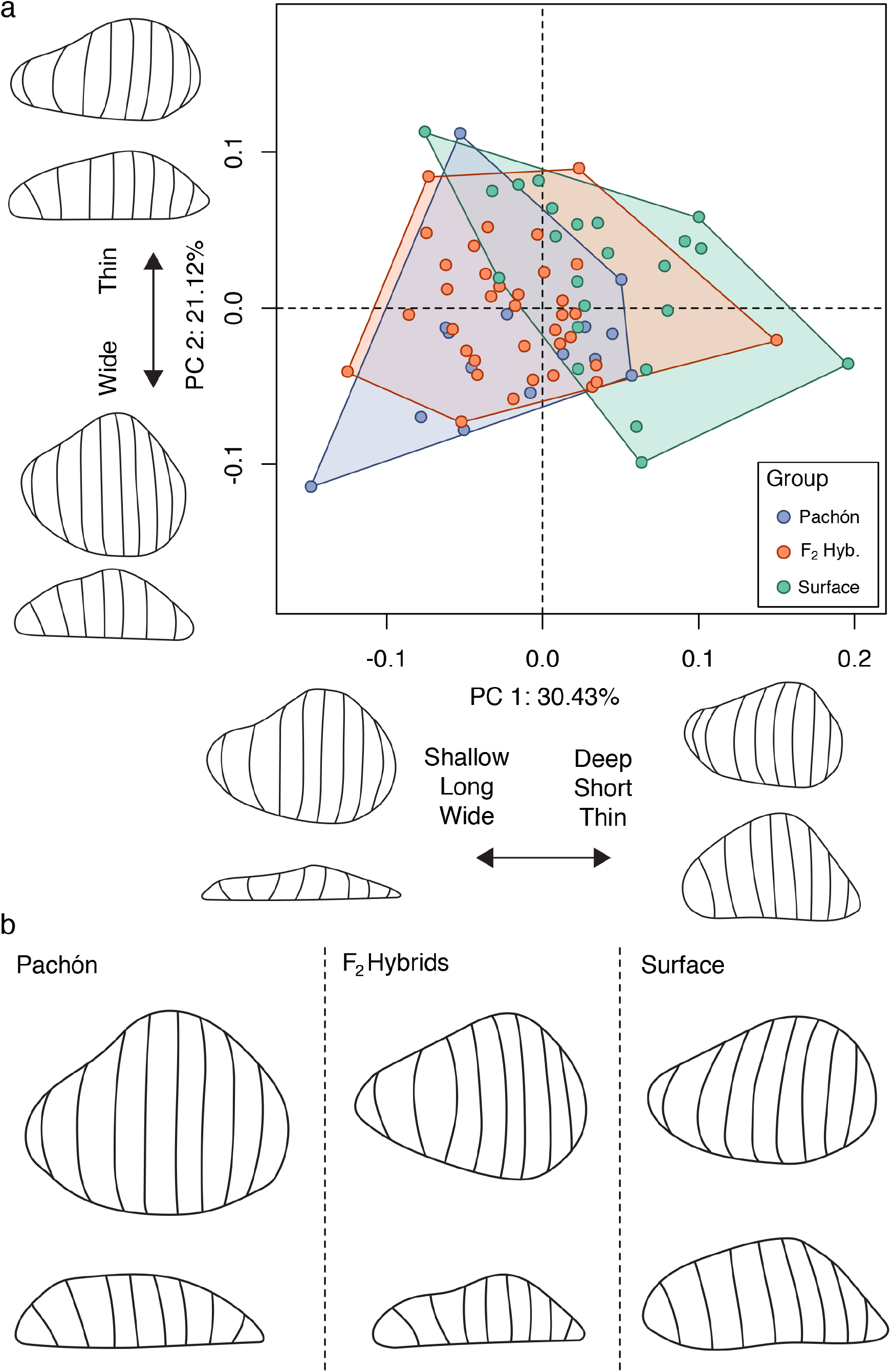
Pineal shape variation in hybrids exhibits a cavefish dominant phenotype. (**a**) PCA capturing 50 percent of the variation across surface, cave and surface to cave F_2_ hybrids. PC1 describes pineal length, while PC2 describes pineal width across populations. (**b**) Illustrations of the median shape for each population.

